# BRN2 suppresses stress-induced hair greying

**DOI:** 10.64898/2026.09.28.754993

**Authors:** Jeremy H Raymond, Madeleine LE Coz, Zackie Aktary, Marie Pouteaux, Michael Hamm, Evelyne Sage, Vincent Favaudon, Valérie Petit, Alfonso Bellacosa, Colin R Goding, Lionel Larue

## Abstract

Adult stem cells maintain tissue homeostasis and regenerative capacity, but mechanisms preserving their integrity under stress remain incompletely understood. Here, we identify BRN2 (POU3F2), a POU-domain transcription factor, as a regulator of melanocyte lineage homeostasis and response to genotoxic stress. Single-cell transcriptomic analyses identified *Brn2* among transcription factors preferentially active in melanocyte stem cells (McSCs), with BRN2 activity declining with age in human hair follicle melanocytic cells. Melanocyte-specific *Brn2* deletion increased melanocyte differentiation and accumulation in the hair bulb, while downregulating DNA-repair and chromosome-maintenance programs. Despite these changes, *Brn2* loss caused neither spontaneous hair greying nor detectable McSC depletion under steady-state conditions. Following ionizing radiation, however, *Brn2*-deficient cells showed altered DNA-damage responses, reduced mitotic signatures, increased p53 and p21, and enhanced melanocytic differentiation. Irradiated *Brn2*-deficient mice developed rapid, dose-dependent, permanent hair depigmentation associated with McSC and melanocyte depletion. Thus, BRN2 links POU-dependent transcriptional control to stem-cell resilience under genotoxic stress, preserving melanocyte regenerative capacity and protecting against irradiation-induced hair greying.

## Introduction

Stem cells play a critical role in tissue homeostasis, enabling the regeneration of damaged tissues, replacement of cells with high turnover, and long-term self-renewal. Failure of these essential functions contributes to aging. Over the past decade, significant advances have been made in understanding the genes that drive stem cell activation and define stem cell niches (Khlusov *et al*, 2025; Zhang & Chen, 2024). However, far less attention has been paid to the genetic programs that maintain stem cell integrity in response to stress. Identifying factors that safeguard stem cells is critical, as inadequate protection can lead to stem cell depletion, tissue dysfunction, and ultimately loss of organ function.

Melanocyte stem cells (McSCs) provide an excellent model to study these processes. McSCs reside in the bulge of the hair follicle mini-organ and are responsible for generating melanocytes that maintain hair pigmentation (Hsu & Fuchs, 2021; Nishimura, 2011). During embryonic development, pigment-producing melanocytes are derived from melanoblasts originating from the neural crest, which contribute to the first round of hair pigmentation (Larue *et al*, 2013; Vandamme & Berx, 2019). Postnatally, McSCs regenerate melanocytes required for hair pigmentation during successive hair cycles. At the start of anagen phases, McSCs proliferate and give rise to transit-amplifying cells (TACs), which migrate to the mature hair follicle bulb and differentiate into melanocytes, transferring pigment to adjacent keratinocytes (Bertrand *et al*, 2020; Nishimura *et al*, 2002). In mice, the first anagen occurs around one week after birth, the second around five weeks, and the third around 14 weeks.

The failure of this pigmentation system during aging is visible as hair graying and eventual whitening. Both genetic and environmental factors contribute to this process. For instance, loss of BCL2 impairs McSC survival, promoting hair greying (Nishimura *et al*, 2002), and deletion of β-catenin in adult McSCs similarly results in hair whitening (Le Coz *et al*, 2021; Rabbani *et al*, 2011; Takeo *et al*, 2016). Environmental stressors, including UV or ionizing radiation, accelerate hair graying through the generation of reactive oxygen species (ROS) that damage and eventually kill hair follicle melanocytes (Inomata *et al*, 2009; Ueno *et al*, 2014) (Mohri *et al*, 2025)

The microphthalmia-associated transcription factor (MITF) is central to the establishment and maintenance of the melanocyte lineage, coordinating cell survival, differentiation, proliferation, senescence, metabolism, and DNA repair (Goding & Arnheiter, 2019). MITF activity is tightly regulated by activators such as SOX10, PAX3, β-catenin/LEF1, and CREB, as well as repressors including DEC1, ALX3, GLI2, FOXD3, and ATF4. One enigmatic transcription factor that can both upregulate and suppress MITF, depending on cellular context, is BRN2 (Goding & Arnheiter, 2019; Goodall *et al*, 2008; Wellbrock *et al*, 2008).

BRN2, a member of the POU family of transcription factors, is expressed in the brain, including hypothalamic cells (Andersen & Rosenfeld, 2001; Fane *et al*, 2019; Nakai *et al*, 1995; Schonemann *et al*, 1995). Its expression is not detected *in vivo* in melanoblasts from E11.5 to E16.5, nor in cultured melanocytes cell lines (Berlin *et al*, 2012a; Goodall *et al*, 2004b), but it is produced in melanoma tumors and melanoma cell lines *in vitro* (Goodall *et al*, 2004a). Brn2 contributes to melanoma initiation, invasion, and proliferation (Goodall *et al*, 2008; Fane *et al*, 2019; Berlin *et al*, 2012a; Eisen *et al*, 1995; Hamm *et al*, 2021; Zeng *et al*, 2018), and can regulate expression of growth arrest and DNA damage-inducible protein GADD45 following UV irradiation (Lefort *et al*, 2001). Recently, BRN2 has also been shown to participate in DNA repair via interaction with Ku70/Ku80 (Herbert *et al*, 2019), suggesting a role beyond transcriptional regulation. These observations raise the possibility that BRN2 contributes to the response of McSCs to genotoxic stress by coupling the expression of DNA-repair genes with lineage regulation.

Despite its established functions in melanoma and DNA repair, it remains unclear whether Brn2 contributes to the response of adult McSCs to genotoxic stress. In this study, we investigated Brn2 expression in the melanocyte lineage *in vivo* and analyzed its function using melanocyte-specific Brn2 knockout mice. We found that Brn2 is expressed in adult McSCs, TACs, and melanocytes, and that loss of Brn2 leads to premature melanocyte differentiation, increased MITF expression and activity, and reduced expression of DNA-repair genes. Importantly, Brn2-deficient McSCs are highly susceptible to ionizing-radiation-induced genotoxic stress, resulting in a stress-associated differentiation phenotype, melanocyte-lineage depletion, and permanent hair depigmentation.

Our results identify BRN2 as an important regulator of the McSC response to ionizing-radiation-induced stress. Under steady state conditions, Brn2 deletion does not cause progressive hair greying or detectable depletion of McSCs; its requirement becomes apparent after genotoxic challenge, when Brn2 deficiency is associated with melanocyte-lineage loss and permanent depigmentation. The phenotype we describe is therefore conditional, reflecting an altered response of damaged stem cells to genotoxic stress rather than a defect in their steady-state maintenance.

## Results

### Brn2 activity is enriched in melanocyte stem cells and declines with aging

Melanocyte stem cells (McSCs) maintain hair follicle pigmentation throughout life, and depletion of this population leads to hair greying and loss of follicular pigmentation. To identify candidate transcriptional regulators of McSC identity, we performed a comprehensive computational reanalysis of the previously published mouse single-cell RNA-sequencing dataset from Berns and colleagues (Berns *et al*, 2024). This reanalysis, rather than generation of the underlying single-cell dataset, is part of the present study and provided the rationale for focusing on Brn2.

Unsupervised clustering allowed for clear distinctions between different cell populations within the mouse hair follicle (Fig. 1A). By evaluating key melanocytic and stem cell markers, this clustering resolved a melanocytic compartment comprising the three established hair follicle melanocyte lineage populations: McSCs in the bulge, transient amplifying cells (TACs) in the intermediate region, and differentiated melanocytes in the bulb (Fig. 1B,C, Appendix Fig. S1A,B).

**Fig. 1.**
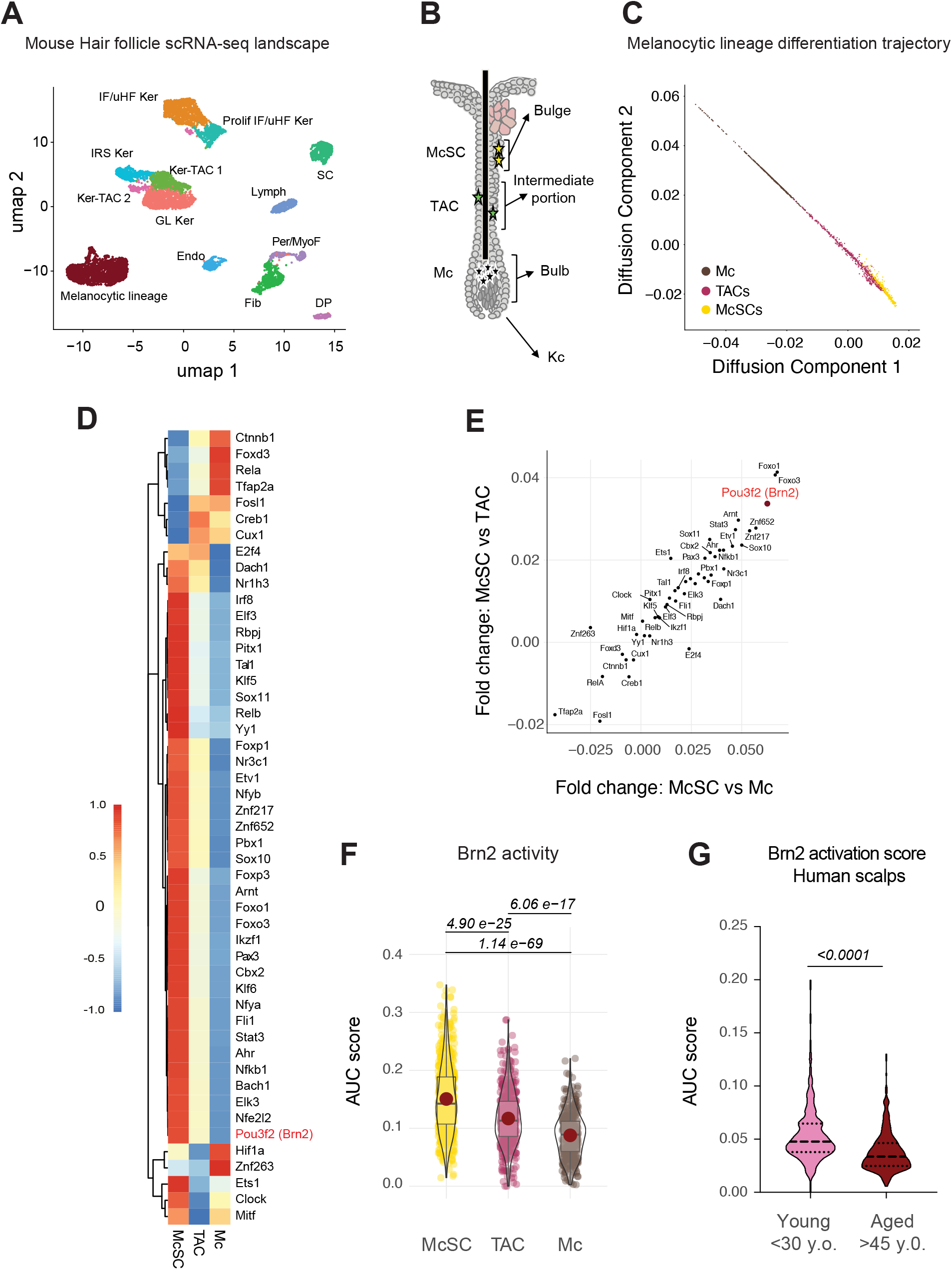
Transcription factor activity in the melanocytic lineage of mouse hair follicles. **A.** UMAP representation of the previously published mouse skin scRNA-seq (Berns et al.) showing major cell types: Endo: Endothelial Cells; Lymph: Lymphatic Vessel Cells; Per/MyoF: Pericytes/Myofibroblasts; Fib: Fibroblasts; DP: Dermal Papilla; SC: Schwann Cells; IFE/uHF Ker: Interfollicular/upper Hair Follicle Keratinocytes; Prolif IFE/uHF Ker: Proliferative Interfollicular/upper Hair Follicle Keratinocytes; GL Ker: Germinative Layer Keratinocytes; Ker-TAC: Keratinocyte Transit Amplifying Cells; IRS Ker: Inner Root Sheath Keratinocytes. **B.** Schematic of the hair follicle depicting the localization of Melanocyte Stem Cells (McSCs), Transit Amplifying Cells (TAC); Melanocytes (Mc), and Keratinocytes (Kc) **C.** Diffusion map of the mouse hair follicle melanocytic lineage **D.** Heatmap of average transcription factor activity for McSCs, Mel-TACs, and Mel. **E.** Correlation of fold changes in transcription factor activity scores between McSC *vs.* TAC and McSC *vs*. Mel; POU3F2 (Brn2) is highlighted. **F.** Brn2 activity score across melanocytic lineage cell types. Statistical analysis: ANOVA with Holm correction. **G.** Brn2 activity in human scalp melanocytic cells according to patient age category. Statistical analysis: ANOVA with Tukey post-test. This analysis shows an age association and does not establish a causal role for Brn2 in physiological hair greying.

To determine which transcription factors were active in each of these populations, we calculated transcription factor activity scores for individual cells using curated target gene sets associated with the melanocytic lineage (Fig. 1D). Comparison of these scores between McSCs, TACs and Mc allowed for the identification of transcription factors preferentially active in McSCs. Among these transcription factors was Brn2 (Pou3f2), which displayed one of the strongest activity enrichments in McSCs compared to other melanocyte lineage cells (Fig. 1D-F). Notably, analysis of an independent human scalp dataset revealed that Brn2 activity was lower in melanocytic cells from older individuals (Fig. 1G). This age-related association prompted us to investigate the functional role of Brn2. However, the absence of spontaneous greying in non-irradiated Brn2-deficient mice indicates that our findings do not establish a causal role for Brn2 in physiological hair aging in mice.

### Brn2 is not expressed in the melanocyte lineage during embryonic development

Previous studies have shown that Brn2 while BRN2 is expressed in cultured melanoblasts or cultured neural crest (Cook *et al*, 2003; Goodall *et al*, 2004a), its expression is absent in melanoblasts at embryonic day 11.5 (E11.5) (Goodall *et al*, 2004a). Whether it is expressed in melanoblasts at other times in development is not known. To examine whether Brn2 is expressed at later stages of melanocyte development, we used a mouse reporter line (Tyr::Cre; Z/EG^f/+^) in which melanoblasts express GFP (Colombo et al., 2012). GFP-positive melanoblasts were isolated from embryos between E14.5 and E18.5, and Brn2 mRNA levels were assessed by RT-qPCR. Brn2 transcripts were not detected at any of these stages (Fig. 2A). Pten served as a positive control for RNA quality and amplification, while cultured melanoblasts (Melb-a) served as a positive control for BRN2 expression.

**Fig. 2.**
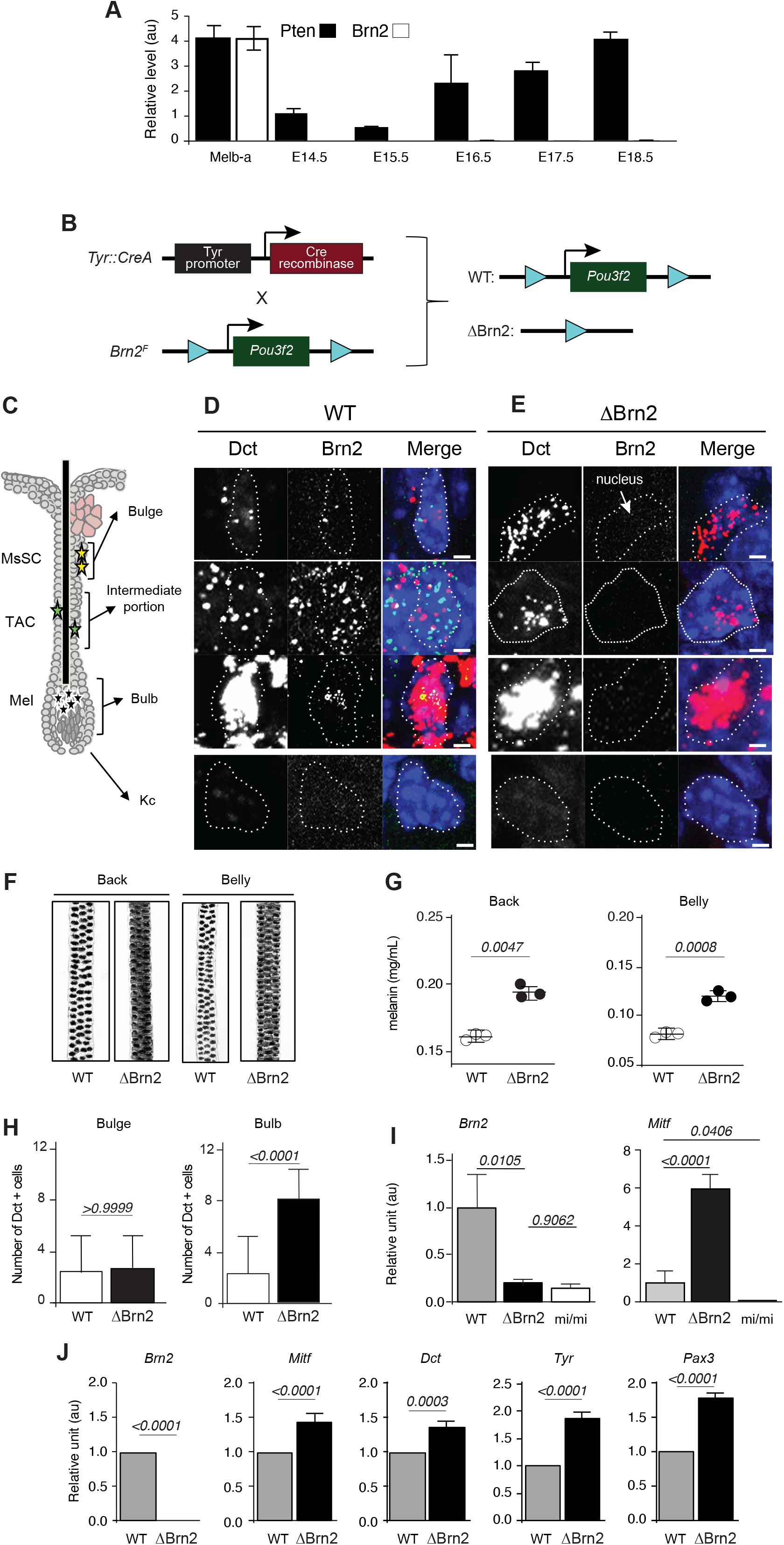
Effects of Brn2 loss in the melanocytic lineage. **A.** RT-qPCR analysis of Brn2 and Pten mRNA levels in GFP+ mouse melanoblasts isolated from embryonic skin during development. Three independent experiments were performed at each stage, with samples analyzed in triplicate. Melb-a cells were used as a cultured melanoblast control. Hprt was used for normalization. **B.** Schematic representation of the mouse model used to generate melanocytic lineage-specific Brn2 deletion (ΔBrn2 = Tyr::Cre; Brn2 ^F/F^, and WT = Brn2^F/F^). **C.** Diagram of an anagen hair follicle showing the anatomical location of melanocytic lineage cells: melanocyte stem cells (McSC) in the bulge, melanocyte transit-amplifying cells (Mel-TAC) in the intermediate region of the follicle, and differentiated melanocytes (Mel) in the bulb. Keratinocytes (Kc) and the sebaceous gland are also indicated. **D. E.** RNAscope detection of Dct and Brn2 mRNA in hair follicle sections from wild-type (WT; Tyr::Cre; Brn2^+/+^) (d) and Brn2-deficient (ΔBrn2; Tyr::Cre; Brn2^F/F^) mice (e). Confocal microscopy images show Dct (red), Brn2 (green), and nuclei (DAPI, blue). Dots correspond to individual mRNA molecules. Dct marks melanocytic lineage cells. The dotted white line delineates a nucleus. Scale bar = 1 μm. **F.** Representative images of hairs from the back and belly of adult WT and ΔBrn2 mice. **G.** Quantification of melanin content in hairs collected from back and belly skin of WT and ΔBrn2 mice. Data are from three independent biological experiments. Statistical significance assessed using an unpaired *t*-test with Welch correction. **H.** Quantification of Dct+ melanocytic lineage cells in the bulge and bulb regions of WT and ΔBrn2 hair follicles. Statistical significance was assessed using an unpaired t-test using data obtained from at least three independent embryos and at least three follicles per sample. **I.** RT-qPCR analysis of Brn2 and Mitf mRNA levels in back skin from postnatal day 4 (P4) WT, ΔBrn2, and melanocyte-deficient Mitf ^vga9/vga9^ (mi/mi) mice. Data are from four independent biological experiments. Statistical significance was assessed using an unpaired *t*-test with Welch correction. **J.** RT-qPCR quantification of melanocytic lineage genes (Mitf, Dct, Tyr, Pax3) in melanocyte cultures derived from WT and ΔBrn2 mice. Gapdh was used as internal control. Error bars represent s.d. Data are from four independent biological experiments. Statistical significance was assessed using an unpaired *t*-test.

These results indicate that, unlike melanoblasts or neural crest in culture, Brn2 is not expressed, or is expressed at levels below detection, in melanoblasts *in vivo* during embryonic development. This suggests that Brn2 may function predominantly postnatally in the melanocyte lineage.

### Brn2 loss in the melanocytic lineage promotes melanocyte accumulation in the hair follicle bulb

Since constitutive *Brn2 (Pou3f2)* knockout mice die within the first 10 days after birth due to severe hypothalamic defects (Nakai *et al*, 1995), we generated mice with a melanocyte-specific deletion of *Pou3f2*, the gene encoding BRN2 (*Tyr::Cre; Brn2^F/F*), to determine the functional role of BRN2 in the melanocyte lineage (Fig. 2B, Appendix Fig. S2A,B). RNAscope analysis confirmed efficient loss of Brn2 expression in melanocytic cells throughout the hair follicle, including the bulge, intermediate region, and bulb (Fig. 2C-E).

Although the overall coat color of mutant mice appeared similar to controls and independent of the number of hair follicle cycles (Appendix Fig. S2C-F), histological analysis revealed a clear increase in melanin deposition within hair follicles lacking Brn2 (Fig. 2F,G). This hyperpigmentation was associated with a marked increase in the number of Dct-positive cells in the bulb, corresponding to differentiated melanocytes, whereas the number of Dct-positive cells in the bulge (McSCs) remained unchanged (Fig. 2H). While surprising, this observation is consistent with human data that showed that human follicles are darker with age in patients not subject to hair greying (Commo *et al*, 2012; Itou *et al*, 2019). Importantly, this increase in differentiated melanocytes was not accompanied by depletion of the stem-cell compartment: the number of Dct-positive cells in the bulge remained unchanged (Fig. 2H), and the coat color of ΔBrn2 mice remained indistinguishable from that of controls throughout life, up to 95 weeks of age, irrespective of the number of hair cycles undergone (Appendix Fig. S2C-F). Because each hair cycle requires renewed McSC activation and production of differentiated melanocytes, progressive exhaustion of the stem-cell pool would be expected to result in a cycle-dependent loss of pigmentation. The absence of such a phenotype indicates that the increased differentiated output associated with Brn2 loss does not compromise long-term maintenance of the McSC compartment under steady-state conditions. Brn2 is therefore dispensable for maintenance of the melanocyte lineage in the unchallenged follicle. Moreover, repeated depilation-induced hair cycles imposed no detectable pigmentary cost in either genotype, indicating that increased proliferative demand alone is insufficient to deplete the McSC pool within the lifespan of a mouse. Thus, replicative challenge and genotoxic stress have distinct consequences for this lineage, and the permanent depigmentation observed following irradiation cannot be explained simply by cumulative proliferative history.

In agreement with this observation, transcriptomic analyses revealed an age-associated increase in the melanocytic gene expression signature in humans (Appendix Fig. S2G). The specificity of Brn2 expression for the melanocyte lineage in the hair follicle was proven by analysing mouse skin lacking melanocytes (*Mitf^mi/mi^ mice*) by RT-qPCR (Fig. 2I). Moreover, RT-qPCR analysis of established melanocytes from ΔBrn2 mice revealed an increased expression of melanocytic genes upon Brn2 deletion (Fig. 2J). Together, these results suggest that Brn2 normally restrains differentiation of the melanocyte stem cell lineage.

### Brn2 deficiency impairs DNA repair pathways in melanocytic cells

To investigate the molecular mechanisms underlying these phenotypes, we performed RNA-seq analysis of control and Brn2-deficient melanocytic cells. Analysis revealed that while the Brn2-low cells increased the level of expression of genes associated with matrix remodeling, their proliferation and DNA repair pathways were downregulated (Fig. 3A). Gene set enrichment analysis (GSEA) revealed a significant downregulation of DNA repair pathways in the absence of Brn2, largely driven by reduced expression of key genes involved in nucleotide excision repair (NER), including *Xpa, Xpc, and Ercc1*, and chromosome maintenance genes such as *Smc2* and *Smc4* (Fig. 3B,C), indicating that Brn2 supports the expression of DNA-repair and chromosome maintenance programs in melanocytic cells. To test whether Brn2-deficient cells display increased sensitivity to genotoxic stress, we examined their transcriptional response following ionizing radiation (IR).

**Fig. 3.**
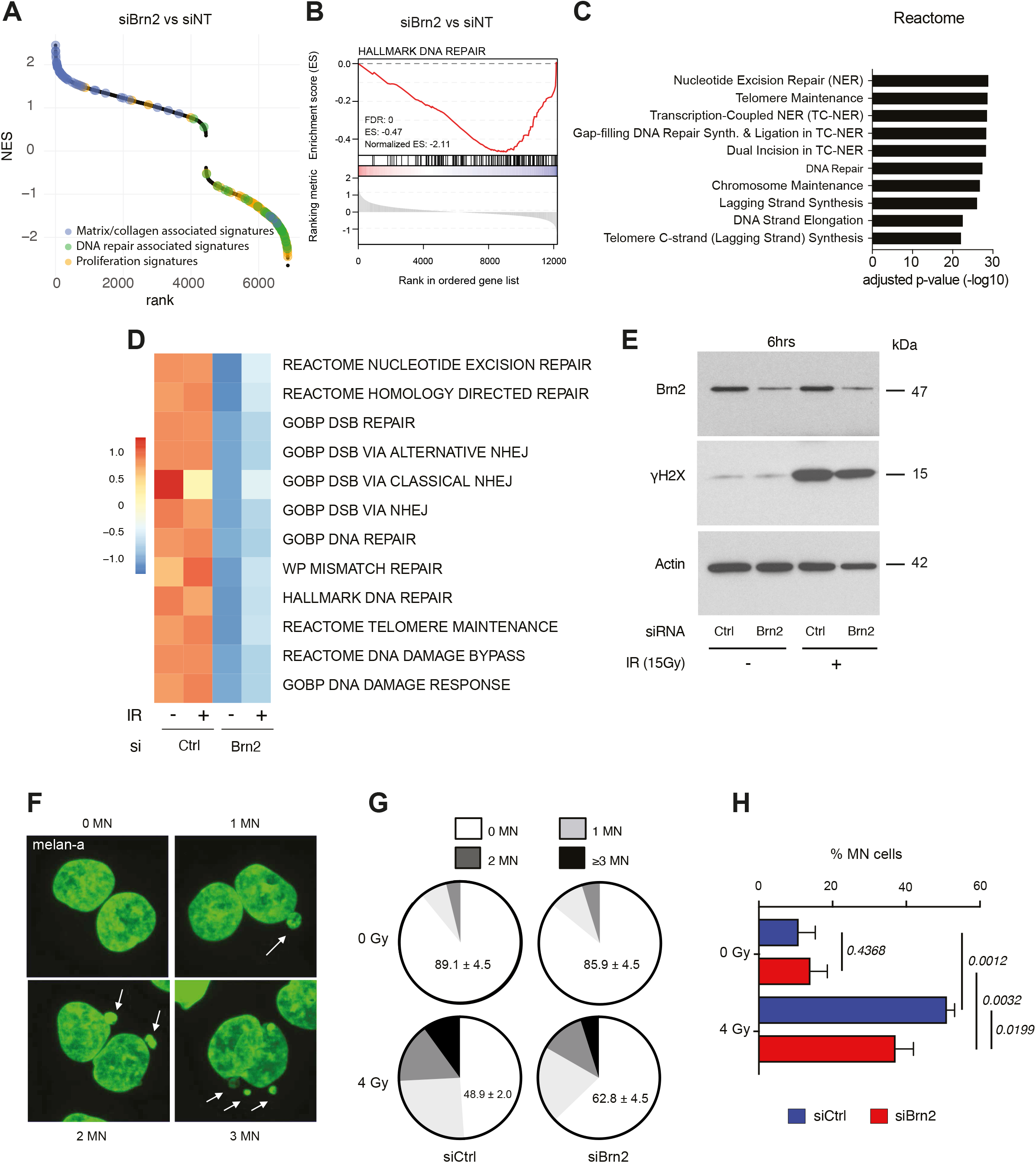
Brn2 regulates DNA repair in melanocytes. **A.** GSEA NES of all MsigDB-C2 gene sets from of siBrn2 versus siNT Melan-a, ranked by NES. Matrix/collagen-associated (blue), DNA repair-associated (green) and proliferation-associated (orange) gene sets are highlighted; positive NES denotes enrichment among genes upregulated after Brn2 knockdown. **B.** Gene Set Enrichment Analysis (GSEA) of DNA-repair signatures in melan-a cells transfected with control siRNA (siCtrl) or Brn2-targeting siRNA (siBrn2). **C.** Reactome enrichment analysis of leading-edge genes from the DNA-repair signature. **D.** Heatmap showing average expression levels of core nucleotide excision repair (NER) and double-strand break (DSB) response/repair genes in melan-a cells following Brn2 knockdown. **E.** Western blot analysis of Brn2 and γ-H2AX protein levels in control and Brn2-depleted melan-a cells after ionizing radiation. Actin was used as a loading control. This blot is interpreted as supportive evidence of altered DNA-damage-response signaling and not as a quantitative measurement of DNA-repair efficiency. **F.** Representative images of nuclei displaying different numbers of micronuclei following irradiation. **G.** Distribution of micronuclei categories in control and Brn2-depleted melan-a cells after irradiation. **H.** Percentage of cells containing micronuclei under control or Brn2 knockdown conditions following irradiation. Statistical significance was assessed using Chi-squared tests with PRISM software based on data from three independent biological experiments.

The loss of Brn2 induced a repression of genes involved in most of the repair pathways independently of IR (Fig. 3D). However, when looking at the core genes of these pathways – such as *Xpa*, *Xpc, Ercc1*, *Ercc2*, *Trp53bp1, or Xrcc4*, IR seems to potentialize the activity of Brn2 (Appendix Fig. S3A). Consistent with these transcriptional alterations, the γH2AX immunoblot showed a lower irradiation-associated signal in Brn2-depleted cells (Fig. 3E). Because this single immunoblot does not by itself provide a quantitative measure of DNA repair efficiency, we now interpret it only as supportive evidence of altered DNA-damage-response signaling and base our conclusion primarily on the transcriptomic DNA-repair signatures. A blunted γH2AX response in cells with elevated MITF is, moreover, the expected outcome rather than an anomaly: upon DNA damage MITF is redirected into a complex with MRE11–RAD50–NBS1 and impairs MRN recruitment to damaged DNA (Binet *et al*, 2024). Because Brn2 represses Mitf^16^, Brn2 loss may therefore attenuate double-strand-break signalling indirectly, through MITF de-repression, in addition to reducing the transcription of nucleotide-excision-repair genes.

Despite the apparent reduction in DNA repair capacity, Brn2-deficient cells exhibited significantly fewer micronuclei following irradiation (Fig. 3F-H; Appendix Fig. S3B-D). Coupled with the decreased proliferative signature observed in melanocyte cells with decreased Brn2 levels (Fig. 3A), these data suggest that cells lacking Brn2 may exit the proliferative cycle rather than accumulate chromosomal instability. Because micronuclei can only be scored in cells that have completed a division, and fewer Brn2-depleted cells remain proliferative following irradiation (Fig. 4D,E), the reduced micronucleus frequency likely reflects, at least in part, the smaller cycling population rather than improved repair of DNA damage. Thus, this finding should not be interpreted as evidence that Brn2 loss enhances genome maintenance.

**Fig. 4.**
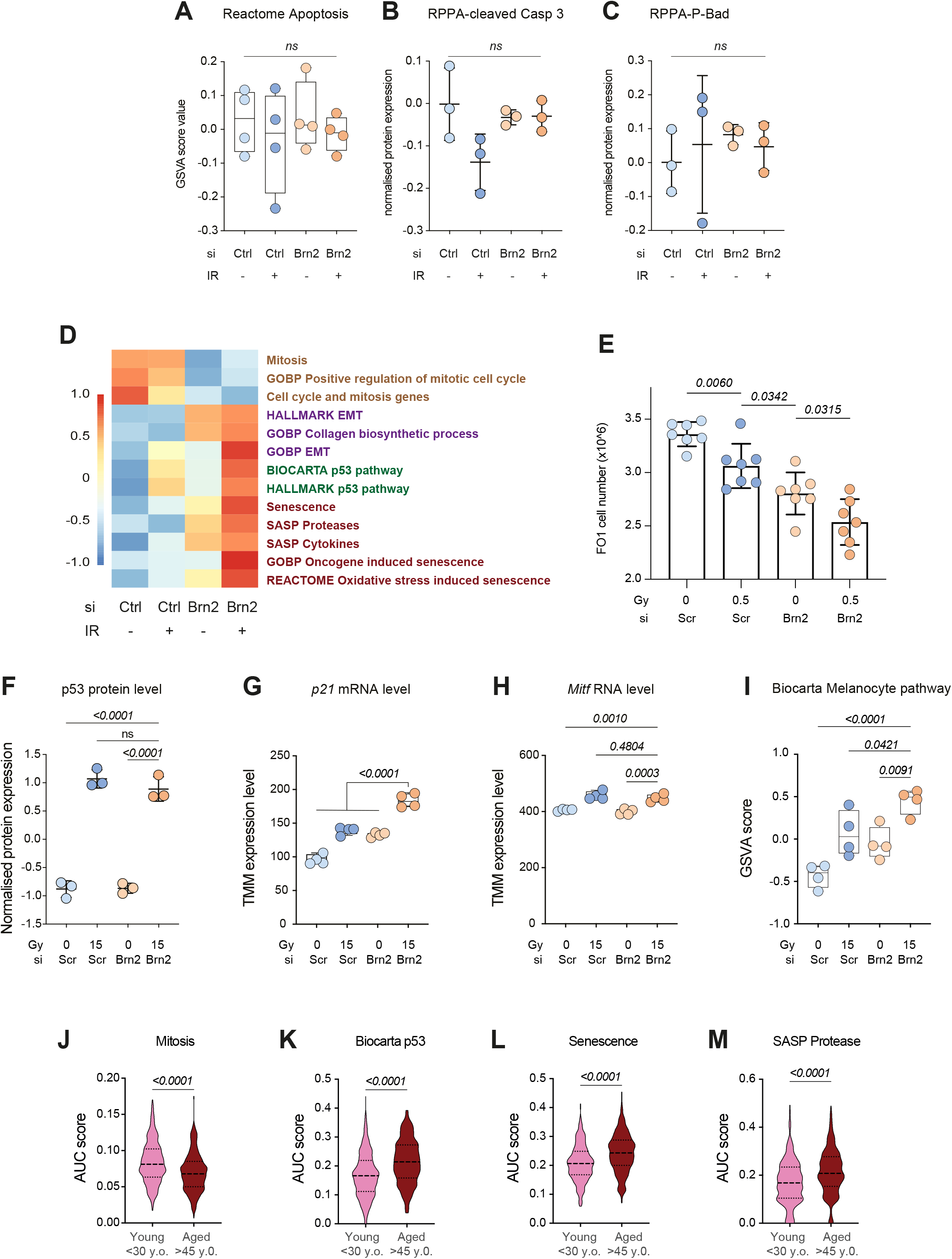
Brn2 modulates apoptosis-associated, mitotic, and stress-induced differentiation in melan-a cells. **A.** Gene Set Variation Analysis (GSVA) scores of the Reactome Apoptosis signature computed from bulk RNA-seq in melan-a cells following Brn2 knockdown and 15 Gy irradiation. **B,C.** Reverse Phase Protein Array (RPPA) quantification of cleaved caspase-3 (**B**) and phosphorylated Bad (**C**) in melan-a cells transfected with control or Brn2 siRNA and exposed to 15 Gy irradiation. **D.** Heatmap of mean GSVA scores for dysregulated signatures computed from bulk RNA-seq in melan-a cells following Brn2 knockdown and 0.5 Gy irradiation. **E.** Quantification of FO-1 cell numbers in cultures with the indicated combinations of Brn2 knockdown and 0.5 Gy irradiation. Statistical significance was assessed using an unpaired t-test based on data from at least six independent biological experiments. **F.** RPPA quantification of p53 protein levels in melan-a cells following Brn2 knockdown and 15 Gy irradiation, showing increased p53 after irradiation and a higher signal in Brn2-depleted irradiated cells. **G,H.** RNA-seq quantification of Cdkn1a (p21; G) and Mitf (H) expression levels in melan-a cells following Brn2 knockdown and irradiation. p21 induction is interpreted as evidence of p53-associated cell cycle control and is not considered a senescence-specific marker. **I.** Biocarta melanocyte pathway GSVA score in melan-a cells following Brn2 knockdown and 15 Gy irradiation. **J-M.** AUC scores for mitosis (J), p53 activation (K), senescence-associated (L) and SASP protease-associated (M) transcriptional signatures in melanocytic cells from human scalp according to patient age category. These are transcriptomic signatures and do not by themselves demonstrate cellular senescence. Statistical analysis: t-test. For panels **A-I** statistical comparisons were performed by Anova followed by a Tukey post-test comparing the siBrn2 + IR group with the other groups, unless otherwise indicated.

### Brn2-deficient cells display a p53-p21-associated stress-induced differentiation phenotype following genotoxic stress

To determine the fate of Brn2-deficient cells after irradiation, we first evaluated whether increased apoptosis accounted for the reduced micronuclei formation. However, analysis at both transcript and protein levels revealed no significant differences in cell death-related gene expression between control and Brn2-deficient cells (Fig. 4A-C).

Instead, RNA-seq analysis revealed a marked reduction in mitotic gene signatures in Brn2-deficient cells (Fig. 4D), indicating decreased proliferative capacity. Furthermore, the number of cells in culture with Brn2 knockdown and irradiation was significantly lower than all other cultures (Fig. 4E).

Ionizing radiation induced robust expression of p53, to a similar extent in control and Brn2-depleted cells (Fig. 4F, Supplementary Table S1). In contrast to p53 itself, the p53 target and cyclin-dependent kinase inhibitor p21 (*CDKN1A*) was significantly upregulated in irradiated Brn2-depleted cells compared with all other conditions (Fig. 4G), consistent with enhanced cell-cycle arrest, although p21 induction alone is not specific for senescence. Importantly, irradiated cells lacking Brn2 expression displayed moderately increased levels of Mitf and a higher melanocyte gene expression signature (Fig. 4H,I). Interestingly, in human scalp melanocytic cells, mitotic gene-expression signatures decreased with age, whereas p53-, senescence- and SASP-associated transcriptional signatures increased (Fig. 4J-M). These transcriptomic associations do not demonstrate bona fide cellular senescence.

Together, these results show that Brn2-deficient melanocytic cells respond to genotoxic stress with reduced proliferation, activation of a p53–p21-associated program, and increased melanocytic differentiation. In the absence of complementary functional senescence assays, we do not conclude that these cells undergo bona fide senescence. Rather, we interpret this phenotype as a stress-induced, senescence-associated differentiation state, reminiscent of the recently described concept of “senodifferentiation” in stem cell systems(Mohri *et al*, 2025). The transcriptional response to irradiation was, moreover, coordinated and extended well beyond p21 induction. In irradiated Brn2-depleted cells, gene sets associated with senescence, SASP cytokines, SASP protease regulators, oncogene-induced and oxidative stress-induced senescence, as well as the BIOCARTA and HALLMARK p53 pathways, all reached their highest enrichment scores, whereas three independent mitotic gene sets reached their lowest (Fig. 4D). Importantly, these gene sets are largely non-overlapping, and neither SASP set contains cell-cycle inhibitors, indicating that this coordinated response cannot simply be attributed to *Cdkn1a* induction. Notably, p53 protein was induced by irradiation to a similar extent in control and Brn2-depleted cells (Fig. 4F), whereas downstream p53 transcriptional activity was more pronounced in the absence of Brn2. Thus, Brn2 loss appears to modify the transcriptional outcome of the p53 response rather than the magnitude of p53 induction itself. Moreover, Brn2 loss alone does not induce this programme but rather enhances the response of the melanocytic lineage once DNA damage has occurred.

These findings should, however, be interpreted in the context of the acute timepoints examined: RNA was collected one hour and protein six hours after irradiation. Since establishment of a bona fide senescent state typically requires a longer period, the absence at these early timepoints of protein-level features associated with established senescence, including changes in p16INK4a (CDKN2A), IL-6 and Rb phosphorylation in our RPPA dataset (Supplementary Table S1), is not unexpected. We therefore interpret these data as evidence for the early induction of a coordinated senescence-associated transcriptional programme rather than as evidence that the cells have already entered an established senescent state.

### Brn2 protects the melanocyte stem cell compartment from irradiation-induced depletion

Given the reduced expression of DNA-repair genes and the altered differentiation responses observed in Brn2-deficient cells, we hypothesized that Brn2 may be required to maintain the melanocyte stem cell pool under genotoxic stress. To test this hypothesis, control and Brn2-deficient mice were exposed to increasing doses of ionizing radiation (10, 15, and 20 Gy) once, at 7 weeks of age, and hair pigmentation was monitored over time.

Whereas control mice maintained largely normal pigmentation following irradiation, Brn2-deficient mice developed pronounced and dose-dependent depigmentation of the irradiated areas (Fig. 5A). Importantly, this depigmentation was permanent over the observation period, consistent with loss of regenerative melanocyte-lineage capacity in the irradiated area. Depigmentation was confined to the irradiated fields, appeared within the first weeks after exposure, and hairs regrowing in these areas remained unpigmented at every subsequent cycle examined (Fig. 5A).

**Fig. 5.**
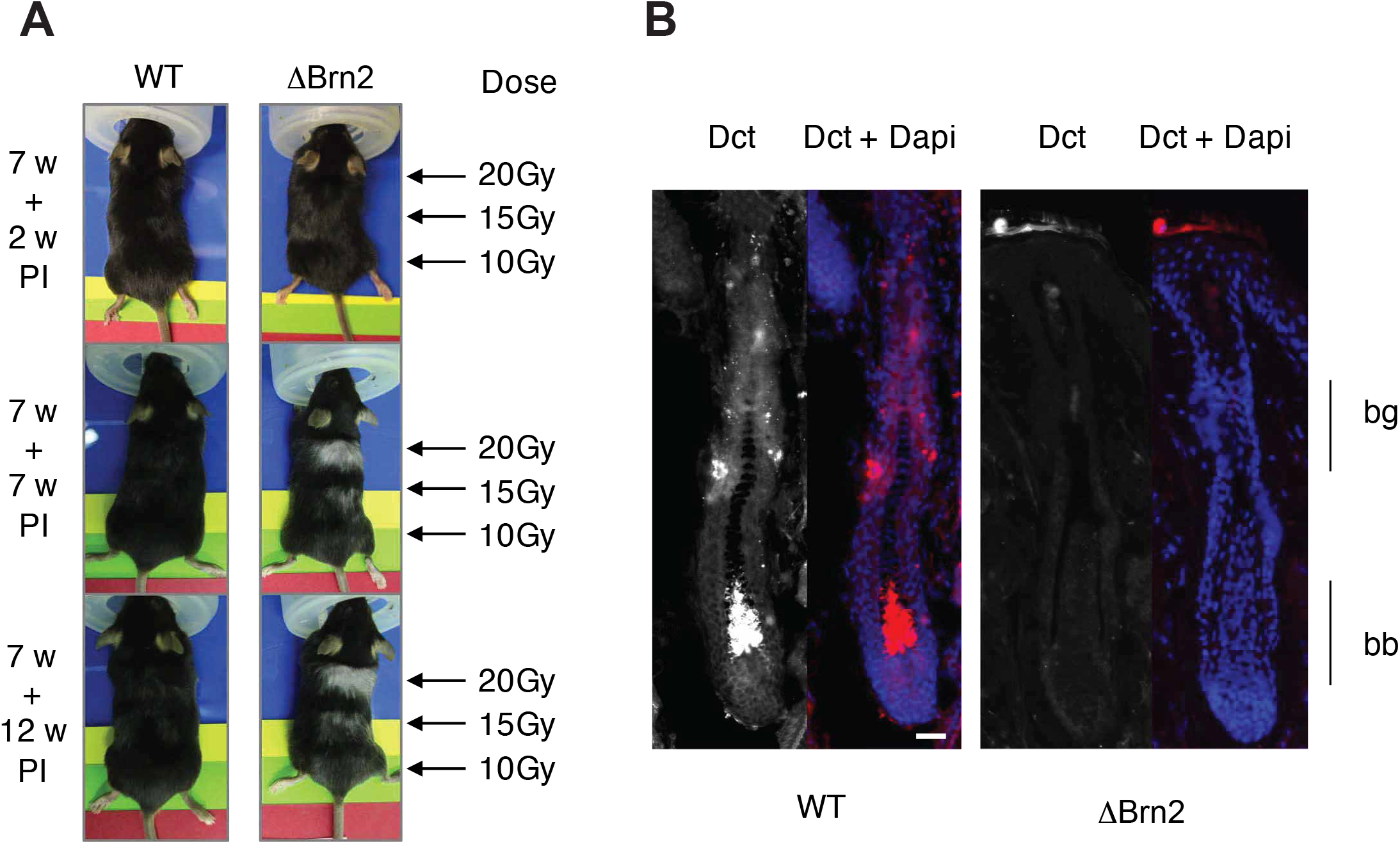
Brn2 deficiency sensitizes hair follicles to irradiation. **A.** Representative photographs of WT and ΔBrn2 mice at 2, 7, and 12 weeks following localized ionizing irradiation. Seven-week-old mice were irradiated on the dorsal skin using a linear accelerator (LINAC) at 10, 15 and 20 Gy. **B.** RNAscope staining for Dct in skin sections from WT and ΔBrn2 mice collected seven weeks after irradiation. Nuclei were counterstained with DAPI. Scale bar = 25 μm. bg = bulge; bb = bulb. The absence of Dct+ cells in irradiated DBrn2 follicles supports loss of the melanocyte lineage after genotoxic stress

Consistent with this interpretation, RNAscope analysis revealed a complete absence of both McSCs and differentiated melanocytes in irradiated hair follicles of Brn2-deficient mice (Fig. 5B), whereas these populations were present in the irradiated hair follicles of control mice. These findings indicate that Brn2 is required for preservation of the melanocyte lineage after ionizing-radiation-induced genotoxic stress.

## Discussion

Hair follicle melanocyte stem cells (McSCs) are essential for maintaining pigmentation throughout life, yet the transcriptional mechanisms that determine their response to genotoxic stress remain poorly defined. Here, we demonstrate that the transcription factor Brn2 (POU3F2) modulates the response of the melanocyte lineage to ionizing-radiation-induced stress, in association with DNA-repair-related transcriptional programs and differentiation control. BRN2 expression is enriched in McSCs and declines with age, and its loss promotes differentiation of McSCs into melanocytes. Furthermore, loss of Brn2 downregulates DNA-repair-associated pathways and alters the response to irradiation, together with reduced proliferation, p53-p21 activation and enhanced differentiation ultimately leading to permanent hair depigmentation following irradiation.

Our findings reveal a dual role for Brn2 in melanocyte lineage regulation. First, Brn2 restrains premature differentiation of McSCs, as its deletion increases the number of differentiated melanocytes in the hair follicle bulb without altering the McSC population in the bulge under steady-state conditions. This is consistent with the elevated expression of MITF and downstream pigmentation genes in Brn2-deficient cells. Second, Brn2 supports expression of DNA-repair-associated pathways, including nucleotide excision repair and double-strand break repair genes. We used the term “DNA-repair-associated” deliberately, as our evidence that Brn2 supports these pathways is primarily transcriptional, whereas the functional consequences of Brn2 loss become apparent only following genotoxic challenge. We do not suggest that Brn2 loss leads to progressive genome instability in the unchallenged follicle; indeed, Brn2 depletion did not increase micronucleus formation under the conditions examined (Fig. 3F-H). Brn2-deficient cells show altered γH2AX signaling and transcriptional responses after ionizing radiation, together with reduced mitotic activity and enhanced differentiation. These observations are consistent with a stress-associated cell-cycle-exit/differentiation response, but do not establish bona fide senescence.

The increased differentiated output at steady state and preservation of the McSC pool are not contradictory. The McSC compartment is dynamic and can compensate for moderate increases in differentiation through self-renewal and progenitor reversion (Sun *et al*, 2023). Our data therefore suggest that Brn2 sets the threshold for McSC differentiation rather than being required for homeostasis: its loss causes increased pigmentation without spontaneous greying, but following genotoxic stress promotes exit from the stem-cell state, leading to rapid and permanent depigmentation. Thus, the requirement for Brn2 becomes apparent under stress rather than at steady state. Although differentiation of damaged melanocytes may transiently contribute to pigmentation, differentiation of McSCs ultimately depletes the reservoir required for subsequent hair cycles. This principle underlies greying induced by irradiation in ATM-deficient McSCs ^12^, senescence-coupled differentiation following DNA damage ^14^, and, as shown here, irradiation in the absence of Brn2. Accordingly, ΔBrn2 follicles show a mild, compensated increase in differentiated melanocytes at steady state, whereas irradiation results in loss of both McSCs and differentiated melanocytes (Fig. 5B), leaving no reservoir from which pigmentation can be regenerated.

The coupling of DNA repair to differentiation control provides a mechanistic explanation for the phenotypes observed *in vivo*. In Brn2-deficient mice, irradiation triggers depletion of McSCs and permanent hair whitening. This in vivo phenotype is consistent with the reduced proliferative and enhanced differentiation programs observed *in vitro*, although the present experiments do not establish that senescence is the causal mechanism. These findings suggest that Brn2 functions as a safeguard, allowing McSCs to maintain proliferative and differentiation potential under stress while preventing premature exhaustion. Importantly, the requirement for Brn2 is stress dependent; non-irradiated Brn2-deficient mice did not show progressive hair greying, and the number of Dct+ cells in the McSC-containing bulge was not decreased under steady state conditions. Thus, our *in vivo* mouse data support a role for Brn2 specifically in preserving the melanocyte lineage in response to genotoxic stress, rather than in physiological aging *per se*.

Notably, while Brn2-deficient cells exhibit defective DNA repair, they paradoxically generate fewer micronuclei following irradiation. This observation likely reflects reduced proliferation of cell-cycle exit rather than reduced DNA damage, highlighting that Brn2 loss shifts the cellular response away from proliferation and toward stress-induced differentiation. We therefore restrict our interpretation to the experimental context examined here: Brn2 influences melanocytic-cell responses to ionizing radiation. Our data do not establish a general mechanism of stem cell aging, nor do they demonstrate that Brn2 protects against apoptosis through a senescence mechanism. Nevertheless, the coordinated changes in DNA-damage response, cell-cycle control, and melanocytic differentiation observed upon Brn2 loss raise the possibility that modulation of the balance between proliferation and lineage differentiation may represent a broader strategy by which stem cells preserve genomic stability and lineage fidelity under conditions of repeated environmental stress or aging. In this context, the stress-induced, senescence-associated differentiation phenotype observed in Brn2-deficient cells may be viewed as reminiscent of “senodifferentiation,” while its functional significance and potential relevance to physiological aging remain to be established.

A stress-dependent requirement has also been reported for another member of the POU family, Oct1/POU2F1, in adult intestinal stem and progenitor cells. Conditional loss of Oct1 had limited effects on intestinal homeostasis under steady-state conditions but impaired epithelial regeneration following DSS-induced injury (Vázquez-Arreguín *et al*, 2019). Although this model involves tissue injury and regenerative stress rather than direct genotoxic stress, and Oct1 and Brn2 belong to distinct POU subclasses, these findings provide an interesting parallel with our observation that Brn2 is largely dispensable for McSC maintenance under steady-state conditions but becomes important following irradiation. Together, these observations raise the possibility that stress-dependent preservation of regenerative capacity may represent a broader, although as yet insufficiently explored, function of POU transcription factors in adult stem-cell compartments.

Interestingly, it was previously shown that Brn2 protects against UVB- and chemotherapy-induced apoptosis by promoting error-prone DNA damage repair in the context of melanoma (Herbert *et al*, 2019). This suggests that the Brn2-mediated protection of McSCs against irradiation-stress revealed here is usurped by melanoma cells to protect against increased double-stranded DNA breaks and replicative stress associated with tumorigenesis (Kloeber & Lou, 2022).

Our study also identifies an association between Brn2 loss, p53-p21 activation, Mitf transcriptional activity, and DNA-repair-associated program after genotoxic stress. Because p53 inhibition was not performed, the present data do not establish that p53 is mechanistically required for the phenotype. Future studies dissecting the direct targets of Brn2 and its interactions with p53 and Mitf will be crucial for understanding the molecular circuitry governing stem cell resilience.

These findings have important implications for hair follicle responses to genotoxic stress. Brn2 loss does not affect hair pigmentation at steady state or during long-term follow up in the absence of irradiation. It is important to note that even prolonged observation in mice is necessarily constrained by their relatively short lifespan and therefore cannot fully recapitulate the decades-long process of physiological hair aging in humans. Conversely, although Brn2 activity is lower in melanocytic cells from older human scalp samples, a direct functional assessment of the consequences of Brn2 loss during physiological aging cannot be experimentally performed in humans. Thus, while the age-associated decrease in BRN2 activity in human melanocytic cells is consistent with a potential contribution to age-related changes in the melanocyte lineage, our data do not establish that this decline is causally responsible for physiological hair greying.

In conclusion, our work identifies Brn2 as an important regulator of melanocyte stem cell homeostasis, linking DNA-repair-associated programs with the control of proliferation and differentiation in response to genotoxic stress. By maintaining this balance, Brn2 contributes to the preservation of the melanocyte lineage following irradiation and limits the stress-induced differentiation response that ultimately leads to permanent hair depigmentation. These findings provide a framework for understanding how transcriptional regulation of the DNA damage response may influence stem cell fate under stress. The relevance of genomic maintenance to tissue homeostasis extends well beyond the melanocyte lineage. Defects in nucleotide excision repair (NER) cause premature aging syndromes, such as *Xeroderma Pigmentosum* and *Cockayne Syndrome* (Bellacosa, 2013; Karikkineth *et al*, 2017; Marteijn *et al*, 2014); whereas increasing evidence links efficient DNA repair and genome maintenance to longevity (Firsanov *et al*, 2025; MacRae *et al*, 2015). In this broader context, our findings raise the possibility that transcriptional coordination of DNA-repair pathways with cell-fate decisions represents an additional layer through which stem cells balance genomic integrity and lineage fidelity during repeated stress. Whether such mechanisms contribute to physiological aging and stem cell exhaustion over the lifespan remains to be determined. More broadly, our findings identify Brn2 as a transcriptional determinant of how damaged melanocyte stem cells respond to genotoxic stress, balancing repair, death and differentiation, with differentiation representing not survival but an irreversible loss of stem-cell potential.

## Materials and Methods

### Mouse hair follicle scRNA-seq analysis

Data from the analysis published by Berns and colleagues (Berns *et al*, 2024) were downloaded from GSE236918. Data were uploaded in R using Seurat (Hao *et al*, 2024) and cells with less than 2^10^ features and more than 20% reads mapping to mitochondrial genes were removed. Doublets were also identified and removed using scDblFinder (Germain *et al*, 2021). Data were integrated based on the first 100 dimensions using harmony. Marker genes were identified using seurat FindAllMarker function and a cell type was attributed to each cluster based on those identified genes. Melanocytic cells were identified based on the expression of Dct. Melanocytic cluster was subset and a diffusion map was created using destiny. Melanocytes were identified by the high expression of differentiated melanocyte genes such as *Oca2*, *Pmel*, and *Mc1r*. McSCs were identified by their highest expression of *Bcl2, Pax3,* or *Klf4*. TACs were identified as the cells in between Melanocytes and McSCs and are characterised by high proliferation scores. Transcription factor activity was computed using the AUCell function of the package escape (Borcherding *et al*, 2021). Transcription factor differential activity was generated using Limma, Data were represented using pheatmap, ggplot2 and ggstaplot (https://doi.org/10.1007/978-3-319-24277-4_9, https://doi.org/10.21105/joss.03167).

### Human scalp scRNA-seq analysis

Data from the analysis published by Ober-Reynolds and colleagues were downloaded from <u>GSE212450</u>. Data were uploaded in R using seurat and cells with less than 300 features and more than 10^4.5^ counts and 25% reads mapping to mitochondrial genes were removed. Data were integrated based on the first 100 dimensions using harmony. Melanocyte cluster was identified based on the expression of DCT. Transcription factor activity was computed using the AUCell function of the package escape.

### Generation of a transcription factor activity signatures in melanocytic cells

To generate an unbiased list of TF targets in melanocytic cells, TPM expression data from The Cancer Genome Atlas TCGA-SKCM were downloaded using TCGAbiolinks (Colaprico *et al*, 2016). The expression of each gene was correlated to all the other ones using pearson. Only genes with a correlation >0.4 with the «prey» gene were kept. Transcription factors (TFs) were subseted from this list and only the genes shown to be bound by the TFs in ChIP-seq analysis were kept. The list of ChIP-seq used was ChEA 2022. The list of transcription factors and their targets is provided in Supplementary Table S2.

### Transgenic mice

Tyr::Cre and Brn2 F/F were previously generated (Delmas et al. 2003; Jaegle et al. 2003). All transgenic mice were maintained on a C57BL6/J background. Animal care and use for this study were performed in accordance with the recommendations of the European Community (2010/63/UE) for the care and use of laboratory animals. Experimental procedures were specifically approved by the ethics committee of the Institut Curie CEEA-IC #118 (CEEA-IC 2016-001) in compliance with the international guidelines.

### Cell lines

Melanocyte cell lines were established in cell culture from the skin of 1.5 day old WT and ΔBrn2 littermate pups as previously described (Berlin *et al*, 2012b). WT, ΔBrn2 and Melan-a mouse melanocyte cells were all grown in F12 medium (Gibco, 21765-029) supplemented with 10% FBS (Gibco, 10270-106), antibiotics (100 U/mL penicillin and 100 μg/mL streptomycin; Gibco, 15140), and 200nM TPA (Sigma, P1585). FO-1 human melanoma cell lines were grown in RPMI 1640 media (GIBCO, 11875101), supplemented with 10% FBS and 1% antibiotics. All cell lines were maintained at 37°C in a humidified atmosphere containing 5% CO_2_.

### BRN2-CRISPR plasmid construction

CRISPR-Cas9-empty plasmids were a generous gift from Thomas Zillinger (University Hospital Bonn, Germany). CRISPR-BRN2-knockout and CRISPR-GFP-knockout plasmids were cloned by linearizing the CRISPR-Cas9-empty plasmid with *SwaI* (NEB, R0604) in NEB buffer 3.1 for 1hour at 25°C, followed by heat inactivation of the enzyme at 65°C for 20 minutes. A target-specific oligo was then annealed with the CRISPR Universal antisense oligo (each 100μM) and denaturated at 95°C for 5 min following a temperature gradient of 0°C/30s down to 8°C. The annealed oligos were filled up with 0.4μL of dNTP’s (10μM) and incubated at 37°C for 1hour followed by an inactivating step of 75°C for 10 min. 0.5μL of the filled in oligos were mixed with 1μL (15ng) of the purified linearized CRISPR-Cas9-empty plasmid, mixed with 1.5μL H_2_O and 2,5μL Gibson Assembly Master-mix (2x Gibson Master Mix [250μL]: 100μL 50% PEG 5000 [w/v], 50μL 1M Tris pH 7.5, 15μL H_2_O, 5μL 1M MgCl_2_, 10μL dNTP’s [10μM], 10μL 50mM NAD^+^ [NEB, B9007S], 5μL 1M DTT, 50μL Taq Ligase [40U/μL; NEB, M0208S], 6.25μL Phusion DNA Polymerase [2U/μL; NEB, MO530S] and 0.2μL T5 exonuclease [10U/μL; NEB, M0363S]) and incubated at 50°C for 1 hour. Two μL of this mix was then used to transform ultracompetent bacteria XL10 Gold (Stratagene, 200314) according to manufacturer’s instructions. The next day, clones were picked and grown over night on a shaker at 37°C in 5 mL LB medium with ampicillin (100 μg/mL). Plasmid DNA isolation was performed according to the manufacturer’s instructions of the Qiagen Quiaprep Spin Miniprep Kit (Qiagen, 27104). Plasmid preparations (1μg) were analyzed by agarose gel electrophoresis (AGE) after digestion with *Swa*I and *EcoR*V (NEB, 3195) in NEB buffer 3.1.

### CRISPR transfection and cell line generation

FO-1 cells were seeded on 10cm cell culture dishes and transfected at 60% confluency with 24μg of the respective CRISPR-Cas9 plasmid per plate using Lipofectamine 2000 (ThermoFisher, 11668019) according to the manufacturer’s protocol. After 48 hours, Blasticidine (4μg/mL) was added to the cell culture media and maintained for three weeks. Blasticidine-resistant cells were then trypsinized, washed and resuspended in PBS for single-cell sorting using the BD FACS ARIA III cell sorter. Single-cell clones were established and propagated in growth media containing Blasticidine (4μg/mL). Isolated clones were validated by western blot for presence/absence of BRN2 protein. Genomic DNA of selected clones was extracted using the AllPrep DNA/RNA Mini Kit (Qiagen, 80204) and subjected to DNA sequencing to determine the introduced mutation.

### RNAseq

For RNAseq experiment, melan-a mouse cells were transfected using lipofectamine 2000 (Invitrogen) with 100pmoL of siRNA targeting mouse BRN2 (Dharmacon M-020029-01-0005) or a Scrambled sequence. Cells were irradiated with GSR D1 irradiator Cs-137 at a dose of 15Gy. Transfections and irradiations were done 4 times on consecutive passages of the cells. RNA was extracted 1 hour after irradiation using the miRNAeasy kit (Qiagen, 217004), according to the manufacturer’s protocol. RNA integrity (RIN) was measured using an Agilent Bioanalyser 2100 and a RNAnano 6000 kit (5067-1511, Agilent Technologies). All RNA had a RIN of 10. RNA concentrations were measured using a NanoDrop (NanoDrop Technologies). RNA-sequencing libraries were prepared from 1μg total RNA using an Illumina TruSeq Stranded mRNA library preparation kit, which allows strand-specific sequencing. PolyA selection using magnetic beads was done to focus the sequencing on polyadenylated transcripts. After fragmentation, cDNA synthesis was done and the resulting fragments used for dA-tailing, followed by ligation with TruSeq indexed adapters. The fragments were amplified by PCR to generate the final barcoded cDNA libraries (12 amplifications cycles). The libraries were equimolarly pooled and subjected to qPCR quantification using the KAPA library quantification kit (Roche). Sequencing was carried out on a NovaSeq 6000 instrument (Illumina) based on a 2 × 100 cycle mode (paired-end reads, 100 bases) using an S1 flow cell. Reads were mapped to the mm10 mouse reference genome (gencode m13 version-GRCm38.p5) using STAR. STAR was also used to create the expression matrices. Reads were mapped to the mm10 mouse reference genome (gencode m13 version-GRCm38.p5) using STAR (Dobin *et al*, 2013). Batches were corrected with Combat (Leek *et al*, 2012). Differential gene-expression analysis was done following the DEseq2 pipeline (Love *et al*, 2014). GSEA was done using fgsea (https://doi.org/10.1101/060012) on signature at least 15 genes. Other signature scores were obtained by GSVA. The list of the signature used for the analyses is provided in Supplementary Table S3. Graphical representations were made with ggplot2 (https://doi.org/10.1007/978-3-319-24277-4) and pheatmap.

### RPPA

For RPPA experiment, melan-a mouse cells were transfected using lipofectamine 2000 (Invitrogen) with 100pmoL of siRNA targeting mouse BRN2 (Dharmacon M-020029-01-0005) or a Scrambled sequence. Cells were irradiated with GSR D1 irradiator Cs-137 at a dose of 15Gy. Transfections and irradiations were done 3 times on consecutive passages of the cells. Total proteins were extracted 6 hours after irradiation with the following lysis buffer : 1% Triton X-100, 50mM HEPES, pH 7.4, 150mM NaCl, 1.5mM MgCl2, 1mM EGTA, 100mM NaF, 10mM Na pyrophosphate, 1mM Na3VO4, 10% glycerol, containing freshly added Complete protease inhibitor (Roche, 11873580001) and PhosStop phosphatase inhibitor cocktail (Roche, 4906845001). Cell lysates were mixed with 4X sample buffer (40% glycerol, 8%SDS, 0.25M Tris-HCL Ph6.8, beta-mercaptoethanol), boiled for 5 minutes and store at −80°C until sample submission to the MD Anderson Cancer Center. Protein extracts were spotted onto nitrocellulose-coated slides arranged in a 4×12-pin-11×11 subarray that accommodate 1056 serially diluted samples plus replicate vertical controls on each slide. Proteins of interest were detected by probing with specific antibodies, amplifying the signal via a tyramide amplification system and visualized via DAB colorimetric reaction. The detection system was a GenPoint-based staining kit from Agilent. Digital images of slides were obtained by scanning on a Huron TissueScope scanner producing 16-bit TIFF files. Spot intensities from the TIFF files were determined via Array-Pro Analyzer software. Mean net intensities of spots from all 1056 samples were used for curve-fitting for each slide (RPPA Super Position and Concentration Evaluation, aka SPACE). Relative protein level for each sample was determined by converging its sample dilution series to one point (EC_50_) and interpolating in RPPASPACE. Expression was normalised sample loading differences by bidirectionally median centering values by antibody then by sample and then adjusted for Set-to-Set differences (batch effects) by normalizing identical control samples in this set with an invariant control sample set and applying the adjustment (differences in means × inverse of standard deviation ratio) to each corresponding data point.

### siRNA transfection and irradiation of cell lines

Melan-a and FO-1 cell lines were seeded on 6-well plates 24-hours prior to transfection. When at 70% confluence, cells were transfected using lipofectamine 2000 (Invitrogen) according to the manufacturer’s instructions with 1μg plasmids or 100pmoL of the scrambled or Brn2-specific siRNAs (Dharmacon). Six hours after transfection, the culture medium was changed.

Cells received 0.5 or 15 Gy γ-rays, depending of the cell type, using a ^137^Cs GSR D1 irradiator (Gamma-Service Medical GmbH) twenty-four hours after transfection. At specific time points post-irradiation, cells were rinsed twice with PBS, and protein or RNA was extracted as described below.

### siRNA sequences

siRNA targeting mouse and human *BRN2* were purchased from Dharmacon as a SMARTpool mix of 4 sequences. The sequences of the siRNA are as follows: for human and mouse *BRN2*: 5’-CGG AUC AAA ACU GGG AUU UA-3’, 5’-GCA AGG GCG CAA GCG GAA A-3’, 5’-AAG GAG GUG GUG AGA GUU U −3’, 5’-CGA CCU UUG CAG GCG AGU AA-3’. A random, scrambled sequence (5’-AAU UCU CCG AAC GUG UCA CGU-3’) was used as a control.

### Semi-quantitative Real-time PCR

Total RNA was extracted from cells using the miRNeasy kit from Qiagen (217004) according to the manufacturer’s protocol. Reverse transcription reactions were performed with 500 ng of total RNA and M-MLV reverse transcriptase (Invitrogen), according to the manufacturer’s instructions. The cDNA generated was then subjected to quantitative real-time PCR for the analysis of gene expression using ABI 7900HT. Oligonulceotides used for PCR are given in Supplementary Table S4.

### Western blot analysis

Whole cell lysates were prepared from melanocyte cell lines using ice-cold RIPA buffer supplemented with complete protease inhibitor cocktail and PhosStop phosphatase inhibitor cocktail (Roche, 11873580001). For western blotting, 25 μg total protein was separated on 15% denaturing acrylamide SDS-PAGE gels and the proteins transferred to nitrocellulose membranes. Membranes were blocked in 5% non-fat milk in Tris-buffered saline supplemented with 0.05% Tween-20 (TBST) and probed with primary antibodies overnight. The signal was detected using peroxidase-conjugated anti-mouse or anti-rabbit secondary antibodies (Jackson 111-035-003 and 111-035-045, respectively) and enhanced chemiluminescence (ECL; Pierce 34075). The primary antibodies used in this study are listed in Supplementary Table S5.

### Epilation of mouse hair

The overall coat color of the mice and color of individual hairs were evaluated as previously described at birth and during the life of adult mice. Hair depilation was performed on mice with a 1:1 mixture of beeswax (Sigma, 243221) and rosin gum (Sigma, R3755). Depilation was first done at seven weeks of age and every four weeks following the first epilation on the back of isofluoran-anesthethized mice. Pictures of the coat color were taken regularly after each epilation.

### Hair count, hair pictures and melanin content assay

Hair plucking was performed on the back and the belly of three mice of each genotype. Random hairs from each genotype were mounted between slides and coverslips in PBS containing 0.1% glycerol and observed under a light microscope. Pictures were taken with light microscope and assembled in Adobe Illustrator. The melanin content of plucked hair follicles was quantified as previously described (Berlin *et al*, 2012b).

### Irradiation of mice and isolation of skin biopsies

Seven week-old mice were irradiated on their backs using a linear accelerator (LINAC) device (Lansonneur *et al*, 2019) with 10, 15 or 20 Gy on different regions of the back. The coat color was documented with pictures over time. Hairs were plucked for observation through the life of the animal and skin biopsies were taken as previously described (Berlin *et al*, 2012b). Mice were euthanized and shaved prior obtaining skin biopsies. Skin was removed and either fixed in 4% PFA in PBS overnight or directly frozen in liquid nitrogen until RNA extraction. The fixed biopsies were included in OCT and frozen over liquid vapours. Cryosections of 10µm thickness were cut and mounted on SuperFrost Plus slides (Fisher Scientific 12-550-15) in the direction of the hair in order to obtain longitudinal sections of the hair follicles. These sections were used for RNAScope staining and immunohistofluorescence.

### RNAscope

Dct mRNA and Brn2 mRNA were detected in skin cryosections of WT and ΔBrn2 mice using RNAscope immunostaining technology following the manufacturer’s recommendations (https://acdbio.com/technical-support/user-manuals) Images were taken using a CLSM SP5 Leica confocal microscope and assembled using ImageJ and Adobe Illustrator.

### Immunofluorescence

Immunostaining of skin sample sections was done using samples fixed in 4% PFA prior to inclusion. All washing steps were done with 0.2% Tween 20 in PBS. Slides carrying skin cryosections were washed and antigene unmasking in citrate buffer was performed while boiling for 20min. The slides were washed again before blocking in PBS + 0.2% Tween 20 and 2% BSA for 1h. The slides were then incubated O/N in a humidified chamber at 4°C with primary antibodies against Dct (Santa Cruz, sc-10451) diluted in PBS + 0.2% Tween 20. The next day, the slides were washed and incubated for 1h at RT in the dark with the secondary antibodies Alexa Fluor 488 (Invitrogen, A21206). The sections were again washed before being incubated with 0.5 µg/µL DAPI to visualize the nucleus. Slides were washed and mounted in Vectashield mounting medium (Vector Laboratories) before imaging by epifluorescence or confocal microscopy. The images were assembled using ImageJ and Adobe Illustrator.

### Micronucleus assay

FO-1 cells were seeded at 1×10^5^ cells per well on glass coverslips in 6 well plates. FO-1 cell lines were transfected with Brn2 or Scrambled siRNAs 24 hours after seeding using Lipofectamine 2000. 24 hours after transfection, the cell lines were irradiated with 0Gy or 4Gy γ-rays from a ^137^Cs irradiator. For the FO-1-CRISPR-GFP and FO-1-CRISPR-BRN2 cells, irradiation was done 48hours after seeding. Immediately after irradiation, the cells were treated with 3μg/mL Cytochalasin B (diluted in fresh growth medium), which was maintained for 30 hours. Glass coverslips were then rinsed twice in PBS and fixed in 4% PFA in PBS 1X for 15 minutes. The coverslips were rinsed again three times in PBS and incubated with DAPI (1:1,000) in PBS for 10 minutes. The glass coverslips were rinsed in PBS three more time before being mounted on glass slides in Vectashield mounting medium. Binucleated cells with or without micronuclei were observed and counted under a fluorescence microscope. At least 500 binucleated cells were counted for each replicate condition. Statistical analysis was performed using Chi-squared tests with PRISM software.

### Statistics

All cell culture experiments were performed with at least three independent biological replicates unless otherwise indicated. RNA-seq experiments were conducted using four independent biological replicates, and RPPA analyses were performed using three independent biological replicates. Data are presented as mean ± standard deviation (SD) unless otherwise specified in the figure legends. Statistical analyses were performed using GraphPad Prism 10 (GraphPad Software) and R (version 4.x). For comparisons between two groups, unpaired two-tailed Student’s *t*-tests with or with Welch’s correction were used. Comparisons involving more than two groups were performed using one-way or two-way ANOVA followed by Tukey’s multiple-comparison test, or using Kruskal–Wallis tests followed by Dunn’s multiple-comparison test when data did not meet parametric assumptions. Differential transcription factor activity analyses from single-cell RNA-sequencing data were performed using Limma, while differential gene expression analyses of bulk RNA-sequencing data were performed using DESeq2 with Benjamini–Hochberg correction for multiple testing. Gene set enrichment analyses were conducted using fgsea, and adjusted *P* values were used to assess significance. Micronucleus frequencies were compared using Chi-square tests. All tests were two-sided unless otherwise stated. Differences were considered statistically significant when *P* < 0.05.

## Acknowledgements

We are grateful to DC Bennett and H Arnheiter for Melb-a cells and mi^vga9^ mice, respectively. Crispr/Cas9 constructs Thomas Zillinger (University Hospital Bonn, Germany). We acknowledge the animal colony (especially Pauline Dubreuil), histology (Sophie Leboucher), imaging (Marie-Noëlle Soler and Claire Lovo) and Christophe Roulin from RadexP facilities of Institut Curie.

## Authors contributions

Jérémy H. RAYMOND conducted the bioinformatics studies, wrote the paper. Madeleine LE COZ conducted experiments and analysis, the mouse work. Zackie AKTARY conducted experiments and analysis. Marie POUTEAUX conducted the mouse work. Michael HAMM conducted experiments and analysis. Evelyne SAGE conducted experiments and analysis. Vincent FAVAUDON conducted experiments and analysis. Valérie PETIT conducted experiments and analysis. Alfonso BELLACOSA interpreted the results. Colin R. GODING interpreted the results. Lionel LARUE designed the study, supervised the overall project, interpreted the results, wrote the paper

## Ethics statement

All methods were performed in accordance with the relevant guidelines and regulations following good scientific practice.

Animal care and use for this study were performed in accordance with the recommendations of the European Community (2010/63/UE) for the care and use of laboratory animals. Experimental procedures were specifically approved by the ethics committee of the Institut Curie CEEA-IC#118 (Authorization #55477-2025040415461498 v2 given by National Authority) in compliance with international guidelines. Mice were housed in a specific-pathogen-free (SPF) certified animal facility with a 12-hour light/dark cycle in a temperature-and humidity-controlled room (22 ± 1 °C and 60%, respectively) with free access to water and food.

## Data availability statement

The RNA-sequencing data generated in this study have been deposited in the Gene Expression Omnibus. The publicly available datasets re-analysed here are available under accessions GSE236918 (mouse hair follicle) and GSE212450 (human scalp). All other data supporting the findings of this study are available from the corresponding author on reasonable request for non-commercial purposes.

## Supplemental information

**Appendix Fig. S1.**
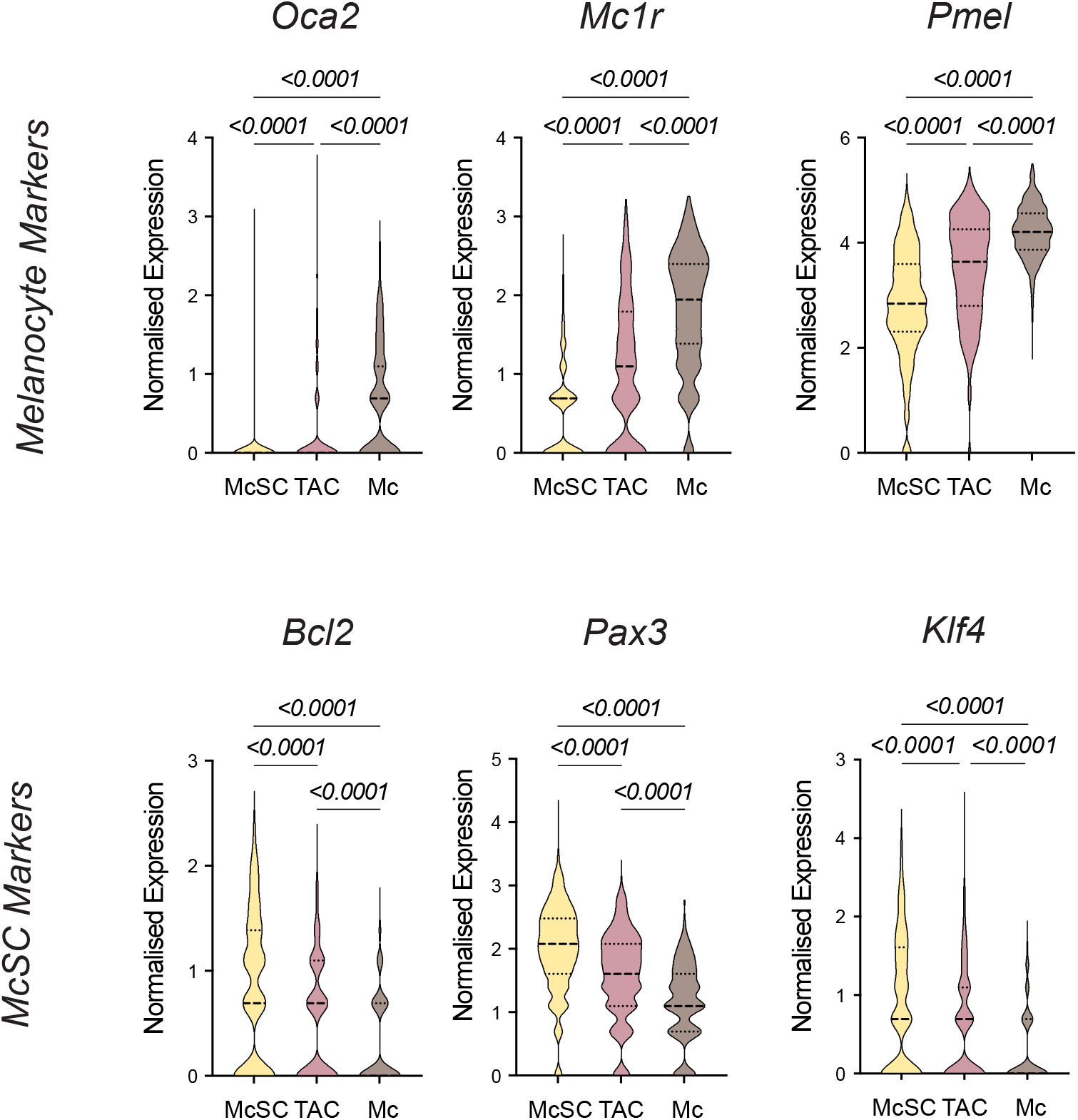
**A.** Expression of some differentiated melanocytes markers – *Oca2*, *Mc1r*, and *Pmel* - in the mouse hair follicle melanocytic cells **B.** Expression of some differentiated melanocytes markers - Bcl2, Pax3, and Klf4 in the mouse hair follicle melanocytic cells In this figure, Data were extracted from Berns et al. McSC: Melanocyte Stem Cells; TAC: Transit Amplifying Cells; Mc: Melanocytes and Kc: Keratinocytes. Statistics were performed by Anova followed by a Tukey’s post test.

**Appendix Fig. S2.**
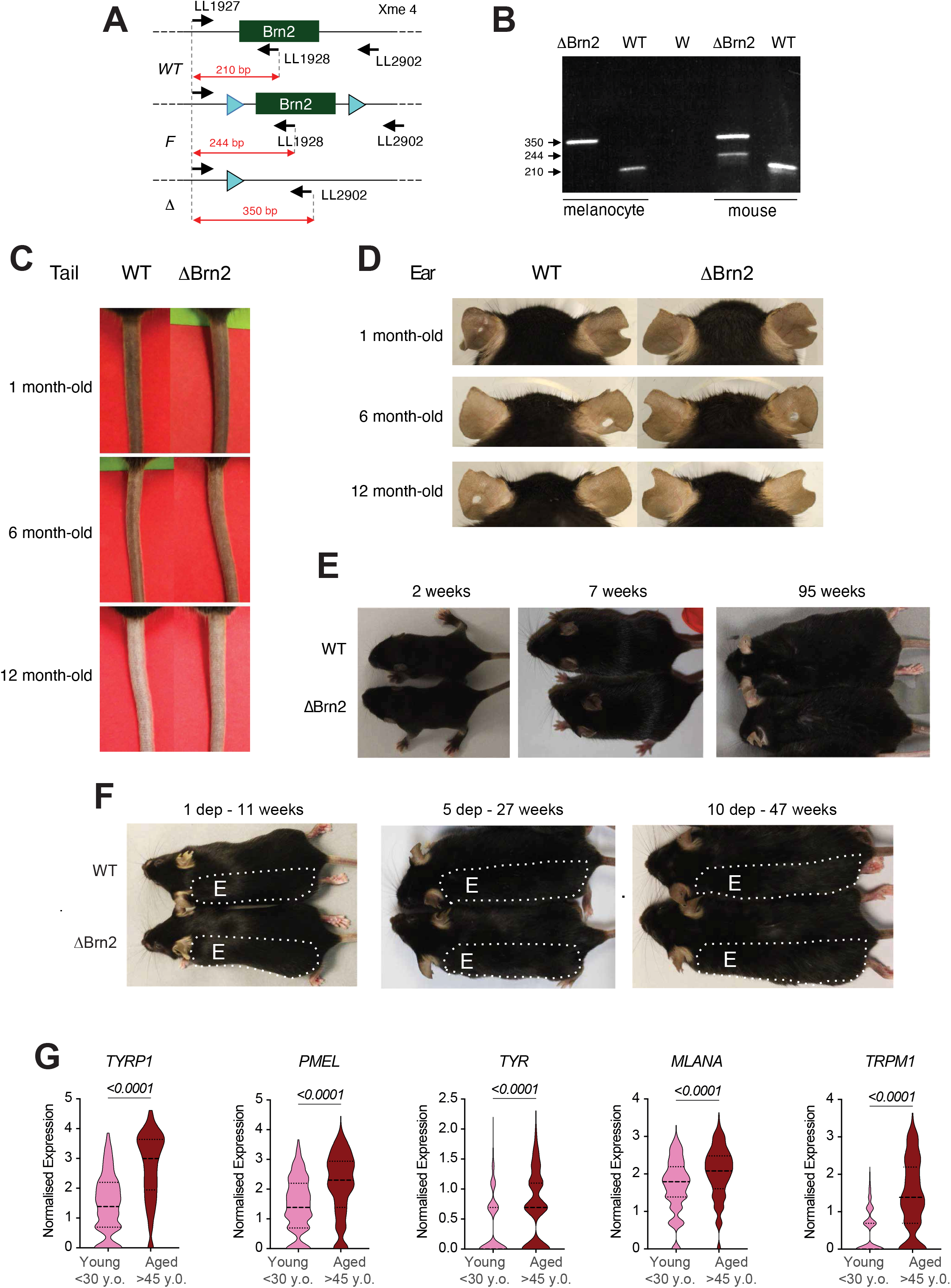
**A.** Genotyping strategy for Pou3f2 in the mouse model. **B.** Representative PCR image for the two main genotypes. In melanocytes and in mouse tails. **C.** Representative image of the tails of WT and ΔBrn2 mice at given time points. **D.** Representative image of the ears of WT and ΔBrn2 mice at given time points. **E.** Representative image of the coat colour of WT and ΔBrn2 mice at given time points. **F.** Representative images of the mice coat colour of WT and ΔBrn2 mice after 1, 5 and 10 depilations. **G.** Expression of the differentiated melanocyte markers TYRP1, PNEL, TYR, MLANA and TRPM1 in the melanocytic cells from human scalps according to patient age.

**Appendix Fig. S3.**
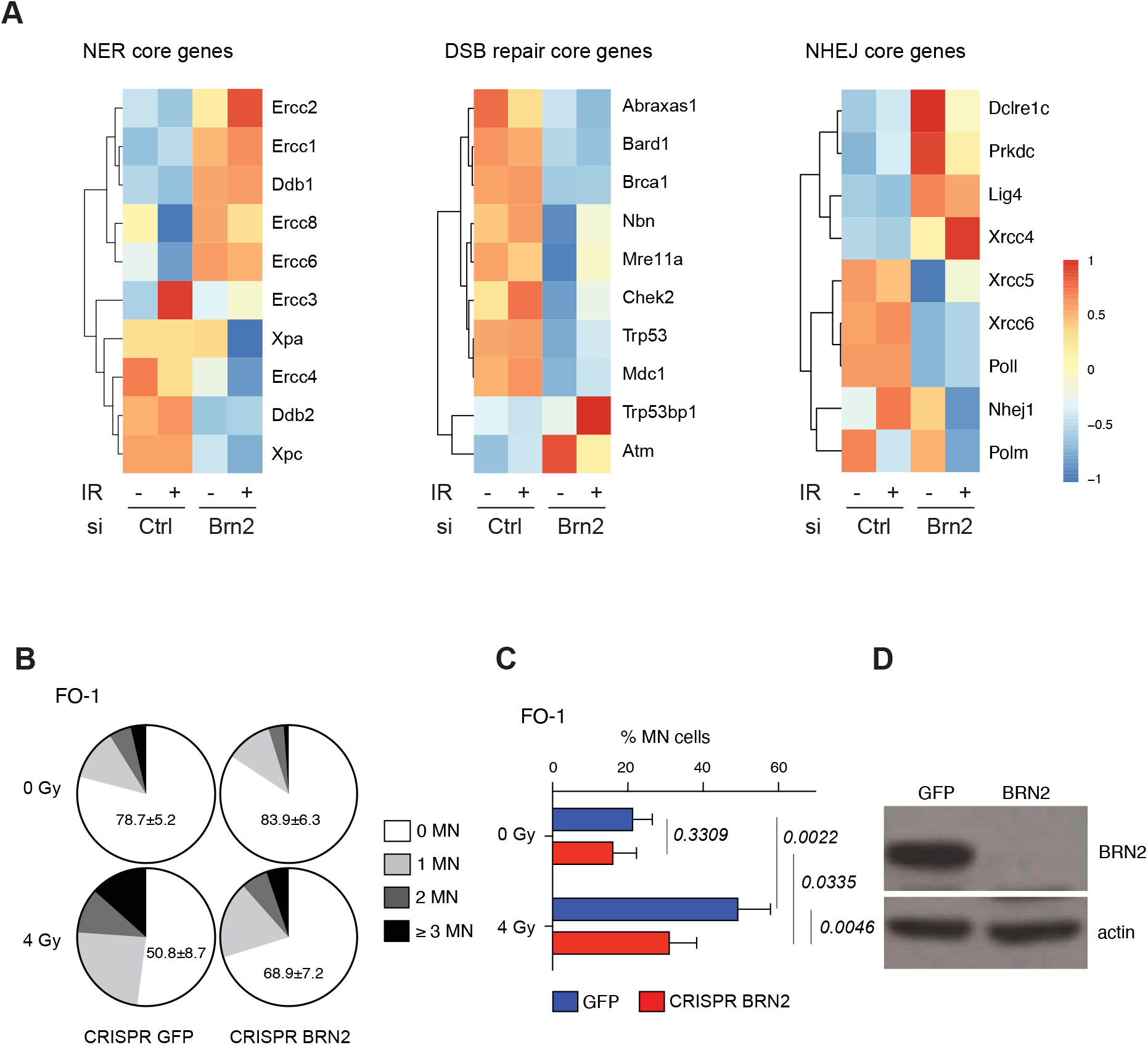
**A.** Heatmap of the average TMM expression of the core gene of the Nucleotide Excision Repair (NER, left), Double Strand Break (DSB, center) and Non-Homologous End-Joining (NHEJ, right) pathways. **B.** Percentage of cells with the given number of micro-nuclei (MN) in FO-1 melanoma cell line K.O. for BRN2 and treated with a 4Gy irradiation. **C.** Quantification of the FO-1 melanoma cell lines harbouring at least 1 MN according to their BRN2 and irradiation status. Statistical significance was assessed using an unpaired *t*-test with Welch’s correction. **D.** representative western blot of the protein expression of BRN2 in the FO-1 melanoma cells K.O. for BRN2.

